# Behavioral flexibility and problem solving in an invasive bird

**DOI:** 10.1101/027706

**Authors:** Corina J. Logan

## Abstract

Behavioral flexibility is considered an important trait for adapting to environmental change, but it is unclear what it is, how it works, and whether it is a problem solving ability. I investigated behavioral flexibility and problem solving abilities experimentally in great-tailed grackles, an invasive species and thus a likely candidate for possessing behavioral flexibility. I found that grackles are behaviorally flexible and good problem solvers, they vary in behavioral flexibility across contexts, flexibility did not correlate with problem solving ability, and those that are more flexible did not necessarily use more learning strategies. It appears that behavioral flexibility can be an independent trait that varies across contexts. Maintaining such a high level of variation could be a mechanism underlying successful species invasions. These results highlight the need to investigate how individuals use behavior to react to changing environments.

## BACKGROUND

Behavioral flexibility, defined here as the ability to change preferences when circumstances change based on learning from previous experience or using causal knowledge, is frequently implicated as a key factor involved in problem solving success and adapting behavior to changing environments (e.g., Lefebvre et al. 1997, Griffin & Guez 2014, Buckner 2015, Chow et al. 2016). Those individuals or species that are more behaviorally flexible are predicted to learn faster and better, and rely on more learning strategies to solve problems (Griffin & Guez 2014). Testing behavioral flexibility experimentally requires individuals to change their behavior in response to changes in the task. Two previous studies investigating behavioral flexibility and problem solving speed found that, contrary to predictions, faster learners were slower to reverse their preferences (invasive Indian mynas: Griffin et al. 2013, threatened Florida scrub-jays: Bebus et al. 2016). Griffin and Guez (2014) propose that behavioral flexibility is a multi-faceted trait: some aspects are measurable in problem solving tasks while other aspects are measurable in other contexts, therefore individuals might exhibit flexibility in some contexts but not others. Behavioral flexibility is usually studied in relation to problem solving speed (Griffin et al. 2013, Bebus et al. 2016), not problem solving success, and it is generally tested only in one context. Therefore, our understanding of the mechanisms underlying behavioral flexibility is lacking.

To begin to address these gaps, I investigated behavioral flexibility in one of the most invasive species in North America, the great-tailed grackle (*Quiscalus mexicanus*, hereafter referred to as grackles; Peer 2011). Species that rapidly adapt to novel environments are presumed to require the ability to behaviorally respond to changing circumstances within the course of their lifetime (Sol & Lefebvre 2000), thus many invasive species are likely candidates for possessing behavioral flexibility. I investigated whether grackles are behaviorally flexible and good problem solvers, whether they vary in behavioral flexibility across contexts, whether flexibility correlates with problem solving ability and speed, and whether individuals that are more flexible use more learning strategies.

I tested behavioral flexibility by measuring initial preferences and then requiring individuals to change preferences after modifying the task in two contexts: a color association task (context 1) and the Aesop’s Fable paradigm (context 2). The color association task (context 1) involved a gold and silver tube placed on the table at the same time and with one of the tubes containing hidden food. Individuals learned to associate food with first the gold tube (learning speed; Experiment 1) and then the silver tube (a modified version of reversal learning; Experiment 6). I used this task to examine which learning strategies grackles used to become proficient. Economics theory predicts solutions to this type of problem, which is called the contextual, binary multi-armed bandit (McInerney 2010). These solutions involve a trade off between an exploration phase and an exploitation phase. The pattern of the trade off determines the learning strategy used.

The Aesop’s Fable paradigm (context 2) examines problem solving ability and involves food floating in a partially filled water tube, which is solved by inserting objects into the tube to raise the water level and bring the food within reach. It has been used to explore the cognitive abilities underlying problem solving in rooks (Bird & Emery 2009), Eurasian jays (Cheke et al. 2011), humans (Cheke et al. 2012), New Caledonian crows (Taylor et al. 2011, Jelbert et al. 2014, Logan et al. 2014), and Western scrub-jays (Logan et al. 2016). While great-tailed grackles are not reported to use tools (Lefebvre et al. 2002), non-tool using species have successfully participated in the Aesop’s Fable tests (Eurasian jays and Western scrub-jays), therefore I expect grackles to be capable of performing these experiments. I compared grackle problem solving performance with previously tested species to determine whether grackles are good problem solvers.

I modified the Aesop’s Fable paradigm to test behavioral flexibility by requiring birds to change preferences using four experiments involving two preference changes, similar to reversal learning experiments, which are considered tests of behavioral flexibility (e.g., Bond et al. 2007, Tebbich et al. 2010, Ghahremani et al. 2010, Buckner 2013). In Experiment 2 (Heavy vs. Light), grackles were given heavy and light objects with the former being twice as functional as the latter, therefore grackles should prefer to insert heavy objects if they attend to the functional properties of the task. However, unlike in most previous experiments (e.g., Cheke et al. 2011, Taylor et al. 2011, Jelbert et al. 2014, Logan et al. 2014, but see Logan et al. 2016), the light objects sank rather than floated, thus if enough were inserted, the food could be reached. I made this modification so that in Experiment 3 (Heavy vs. Light Magic) when the heavy objects became non-functional by sticking to a magnet placed inside the tube above the water, the light objects would now be the functional option because they could fall past the magnet into the water. Individuals that prefer heavy objects or have no preference in the Heavy vs. Light experiment should change their preference in the Heavy vs. Light Magic experiment to preferring neither object or light objects. This would indicate that their preferences are sensitive to changing contexts. Experiments 4 and 5 followed the same methods used for New Caledonian crows (Logan et al. 2014). To solve Experiment 4 (Narrow vs. Wide equal water levels), objects must be inserted into a narrow (functional) rather than a wide (non-functional) tube when water levels are equal in both tubes. In Experiment 5, the narrow tube becomes non-functional because the water level is too low, therefore birds must change their preference to inserting objects into the functional wide tube or to having no preference (as long as they are successful in most trials) to demonstrate behavioral flexibility.

## METHODS

### Ethics

This research was carried out in accordance with permits from the U.S. Fish and Wildlife Service (scientific collecting permit number MB76700A), California Department of Fish and Wildlife (scientific collecting permit number SC-12306), U.S. Geological Survey Bird Banding Laboratory (federal bird banding permit number 23872), and the Institutional Animal Care and Use Committee at the University of California Santa Barbara (IACUC protocol number 860 and 860.1).

### Subjects and Study Site

Eight wild adult great-tailed grackles (4 females and 4 males) were caught using a walk-in baited trap measuring 0.61m high by 0.61m wide by 1.22m long (design from Overington et al. 2011). Birds were caught (and tested) in two batches: batch 1 at the Andree Clark Bird Refuge (4 birds [Tequila, Margarita, Cerveza, Michelada] in September 2014, released in December) and batch 2 at East Beach Park (4 birds [Refresco, Horchata, Batido, Jugo] in January 2015, released in March) in Santa Barbara, California. They were housed individually in aviaries measuring 183cm high by 119cm wide by 236cm long at the University of California Santa Barbara for 2-3 months while participating in the experiments in this study. Grackles were given water *ad libitum* and unrestricted amounts of food (Mazuri Small Bird Food) for at least 20 hrs per day, with their main diet being removed for up to 4 hrs on testing days while they participated in experiments and received peanuts or bread when successful. Grackles were aged by plumage and eye color and sexed by plumage and weight following Pyle (2001). Biometrics, blood, and feathers were collected at the beginning and end of their time in the aviary. Their weights were measured at least once per month, first at the time of trapping using a balancing scale, and subsequently by placing a kitchen scale covered with food in their aviary and recording their weight when they jumped onto the scale to eat.

### Experimental Set Up

Apparatuses were placed on top of rolling tables (60cm wide by 39cm long) and rolled into each individual’s aviary for testing sessions, which lasted up to approximately 20min. If habituation to an apparatus was needed, it was placed in their aviary overnight and they were fed off of it. If an apparatus had parts that would allow a bird to learn how the task worked, these parts were taped over to prevent learning. If a grackle approached an apparatus and ate off it without hesitating, it was considered habituated. If re-habituation was needed, the habituation process was repeated. Color tubes were baited with peanut pieces and/or bread. Water tubes were baited with 1/16 of a peanut attached to a small piece of cork with a tie wrap for buoyancy (hereafter referred to as a peanut float). The area around the top of the tube (the standing platform) was also sometimes baited with smaller peanut pieces and bread crumbs, and more peanut floats could be added to the inside of the water tube to encourage the bird to interact with the task. If more than one peanut float was in the tube, the bird was given the opportunity, after retrieving the first peanut float, to insert more objects into the tube to retrieve the other peanut floats. If a bird started to lose motivation for participating in a task because they were unsuccessful (as in Heavy vs. Light Magic), I baited the standing platform between trials to reward their participation and keep them interested in finishing the experiment. A trial was terminated when the bird solved the task or did not interact with the apparatus. All water tube experiments (2-5) consisted of 20 trials per bird and were recorded with a Nikon D5100 camera on a tripod placed inside the aviary. Experiments are presented in the order they were given, which was the same for all birds. Grackles took 1-7 days to complete an experiment, which could have spanned the course of up to 19 days.

### Experiment 1: Color Association Task (learning speed)

To assess how many trials it takes a grackle to form an association between food and color, they were given a gold and a silver tube with food (peanut pieces or bread) always hidden in the gold tube (Logan et al. 2014 & 2016). Grackles were first trained on a blue tube where they learned to search for hidden food. Each color tube set up consisted of a PVC tube (outer diameter 26mm, inner diameter 19mm) mounted on two pieces of plywood glued together at a right angle (whole apparatus measuring 50mm wide by 50mm tall by 67mm deep. Each tube was placed at opposite ends of a table with the tube openings facing the side walls so the bird could not see which tube contained the food. Tubes were pseudorandomized for side and the left tube was always placed first, followed by the right to avoid behavioral cueing. Pseudorandomization consisted of alternating location for the first two trials of a session and then keeping the same color on the same side for at most 2 consecutive trials thereafter. Each trial consisted of placing the tubes on the table, and then the bird had the opportunity to choose one tube by looking into it (and eating from it if it chose the gold tube). Once the bird chose, the trial ended by removing the tubes.

### Spontaneous Stone Dropping

Birds were given two sequential 5 min trials with the stone dropping training apparatus and two stones to see whether they would spontaneously drop stones down tubes. The stone dropping training apparatus was a clear acrylic box with a tube on top. The box contained out of reach food on to pofa platform that was obtainable by dropping a stone into the top of the tube, which, when contacting the platform, forced the magnet holding it up to release the platform (design as in Bird and Emery 2009 with the following tube dimensions: 90mm tall, outer diameter=50mm, inner diameter=37 or 44mm). The food then fell from the platform to the table. At the end of the first 5 min trial, the stones were moved to different locations on the table and on the wooden blocks. The blocks made it easier to access the top of the tube.

### Stone Dropping Training

Those birds that did not spontaneously drop stones down the tube on the stone dropping training apparatus were trained to push or drop stones down tubes using this same apparatus (Figure 1). Birds were given two stones and went from accidentally dropping stones down the tube as they pulled at food under the stones, which were balanced on the edge of the tube opening, to pushing or dropping stones into the tube from anywhere near the apparatus. Once the bird proficiently pushed or dropped stones into the apparatus 30 times, they moved onto the reachable distance on a water tube. Stone pushing/dropping proficiency was defined as consistently directing the stone to tube opening from anywhere on the ramp on the top of the apparatus. Not all motions had to be in the direction of the tube opening because some grackles preferred to move the stone to a particular location on the ramp (which may initially be in the opposite direction from the tube) and push or drop it in from there or push the stone in shorter, angular strokes. It was permissible for a bird to throw one of the stones off the side of the apparatus (which occurred sporadically throughout all of their experiences with stone pushing/dropping) as long as they proficiently put the other stone in the tube. Similar to Western scrub-jays (Logan et al. 2016), the grackles inserted objects while standing at the top of the tube rather than standing on the ground. The different standing position should not influence their perception of the objects as they were inserted into the tube because their heads were always over the top of the tube at the time of insertion, regardless of where they were standing.

**Figure 1.**
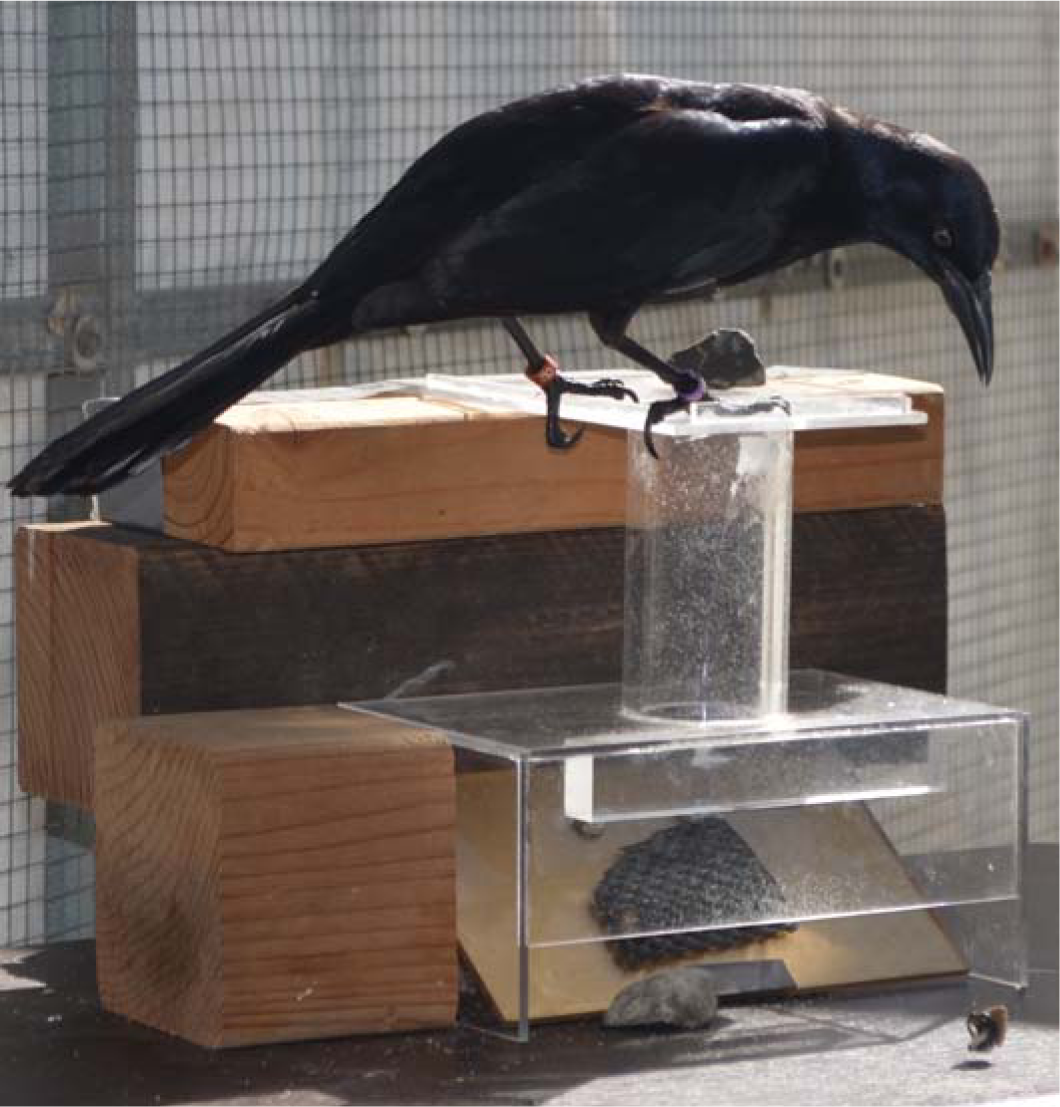
Stone dropping training using a platform apparatus: Batido inserted a stone into the stone dropping training apparatus, which collapsed the platform and the peanut float fell out onto the table.

### Reachable Distance

To determine how high to set the water levels in water displacement experiments, a bird’s reachable distance was obtained. Food was placed on cotton inside a resealable plastic bag, which was stuffed inside the standard water tube (a clear acrylic tube [170mm tall, outer diameter=51mm, inner diameter=38mm] super glued to a clear acrylic base [300×300×3mm]) to obtain the reachable distance without giving the bird experience with water. The food was first placed within reach and then lowered into the tube in 1cm increments until the bird could not reach it. The lowest height the bird could still reach was considered its reachable distance and water levels in subsequent experiments were set to allow the desired number of objects to bring the food within reach.

### Water Tube Proficiency Assessment

To determine whether individuals transferred their stone pushing/dropping skills from a tube on a platform to a tube containing water or whether they needed training on this new apparatus, they were given a standard tube partially filled with water with a peanut float and four stones (9-14g, each displaces 5-6mm water) which they could drop into the tube to raise the water level and consequently reach the food. Once a bird accomplished 30 consecutive proficient trials, they moved onto experiment 1. Proficiency was defined as in the stone dropping training section above.

### Experiment 2: Heavy vs. Light

One standard water tube was presented with 4 heavy (steel rod wrapped in fimo clay, weight=10g,each displaces 2-3mm of water) and 4 light (plastic tube partially filled with fimo clay, weight=2g, each displaces 1-1.5mm of water) objects placed in pseudorandomized (as explained for color learning) pairs near the top of the tube (both objects were 21-24mm long and 8mm in diameter; Figure 2A). Heavy objects had a larger volume (1,056-1,207mm^3^) and displaced 0.5-2mm more water than light objects (volume roughly 500mm^3^), which had a hollow end. Thus the heavy objects were more functional than the light objects, but importantly, both objects were functional. Each bird had three opportunities to interact with the objects before the experiment began: one heavy and one light object was placed on the table (pseudorandomized for side) with food underneath and on top of each object. The object that was first touched was recorded and a trial continued until the bird interacted with both objects. If one object was preferred (as indicated by approaching it first 2-3 times), then more food was placed on the other object to try to eliminate any object preference before the experiment began. Four interactions were given Horchata and 5 to Batido to ensure a lack of preference. After object interaction trials, each bird was given the 20 trial experiment.

**Figure 2.**
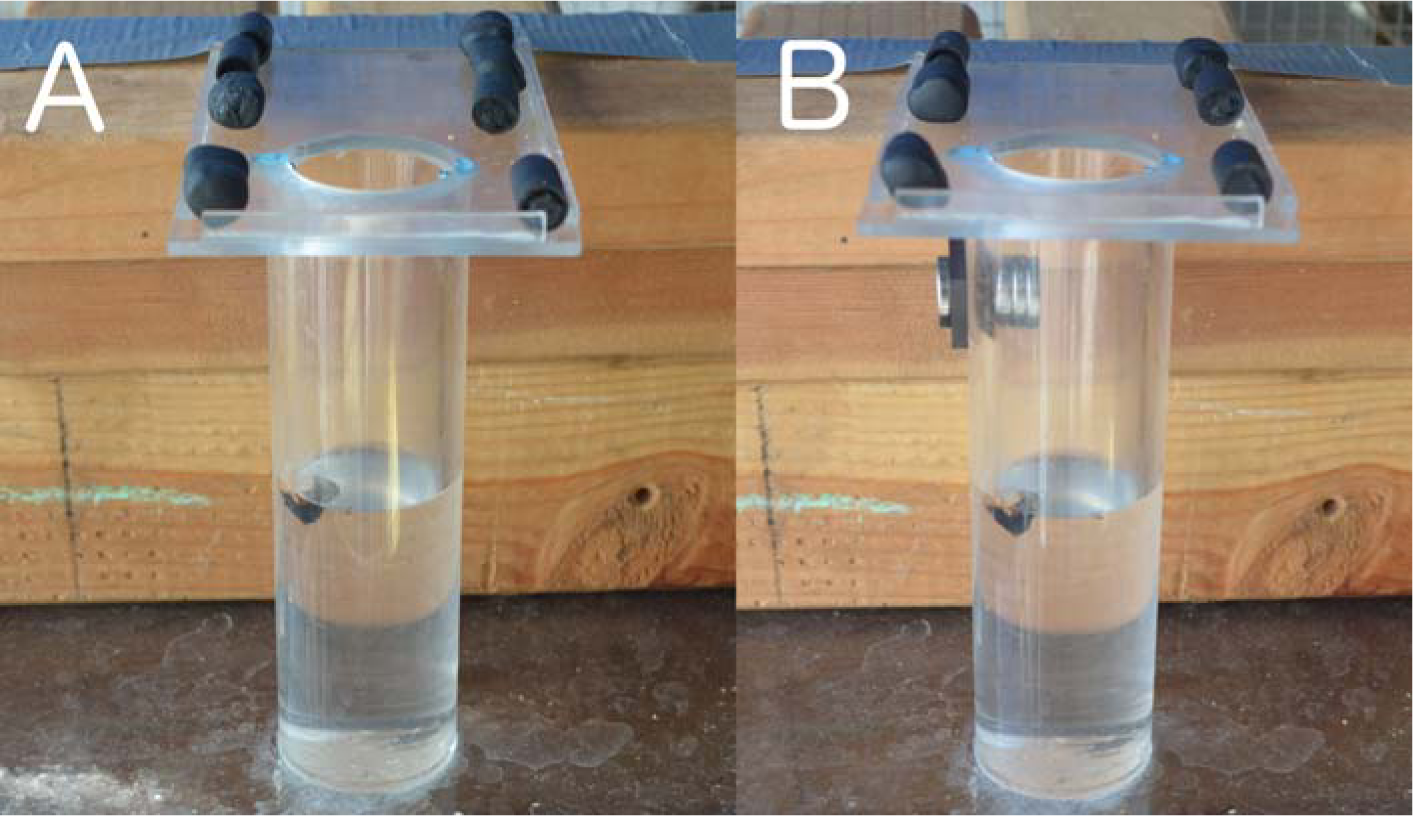
Heavy objects are more functional than light objects in the Heavy vs. Light experiment(A), while the light objects are the only functional objects in the Heavy vs. Light Magic experiment (B).

### Experiment 3: Heavy vs. Light Magic

The set up was the same as in Experiment 1, except there were magnets (2 super magnets on the outside and 3 on the inside of the tube) attached to the tube above the water level such that the heavy objects would stick to the magnets and not displace water, while the light objects could fall past the magnets into the water, thus being the functional choice (Figure 2B). Birds were given 3 heavy and 3 light objects, placed in pseudorandomized pairs near the top of the tube, and 20 trials were conducted.

### Experiment 4: Narrow vs. Wide Equal Water Levels

To determine whether birds understand volume differences, a wide and narrow tube with equal water levels were presented with four objects made out of fimo clay (30×10×5mm, 3-4g, each object displaced 1-2mm in wide tube and 5-6mm in narrow; Logan et al. 2014; Figure 3). Two objects were placed near the narrow tube opening and two objects near the wide tube opening. The objects were only functional if dropped into the narrow tube because the water levels were set such that dropping all of the objects into the wide tube would not bring the floating food within reach. However, dropping 1-2 objects into the narrow tube would raise the water level enough to reach the food. Both tubes were 170mm tall with 3mm thick lids that constricted the opening to 25mm in diameter to equalise the bird’s access to the inside of each tube, and super glued to a clear acrylic base (300×300×3mm). The wide tube (outer diameter=57mm, inner diameter=48mm, volume=307,625mm^3^) was roughly equally larger than the standard water tube (dimensions above, volume=192,800mm^3^) as the narrow tube was smaller (outer diameter=38mm, inner diameter=25mm, volume=83,449mm^3^). The position of the tubes was pseudorandomized for side to ensure that tube choices were not based on a side bias, and 20 trials were conducted. Before the experiment began, each bird had three opportunities to interact with the object, as in Experiment 1, only here it was simply to habituate them to the clay object (one object type) and not to train the birds not to prefer one object type over another.

**Figure 3.**
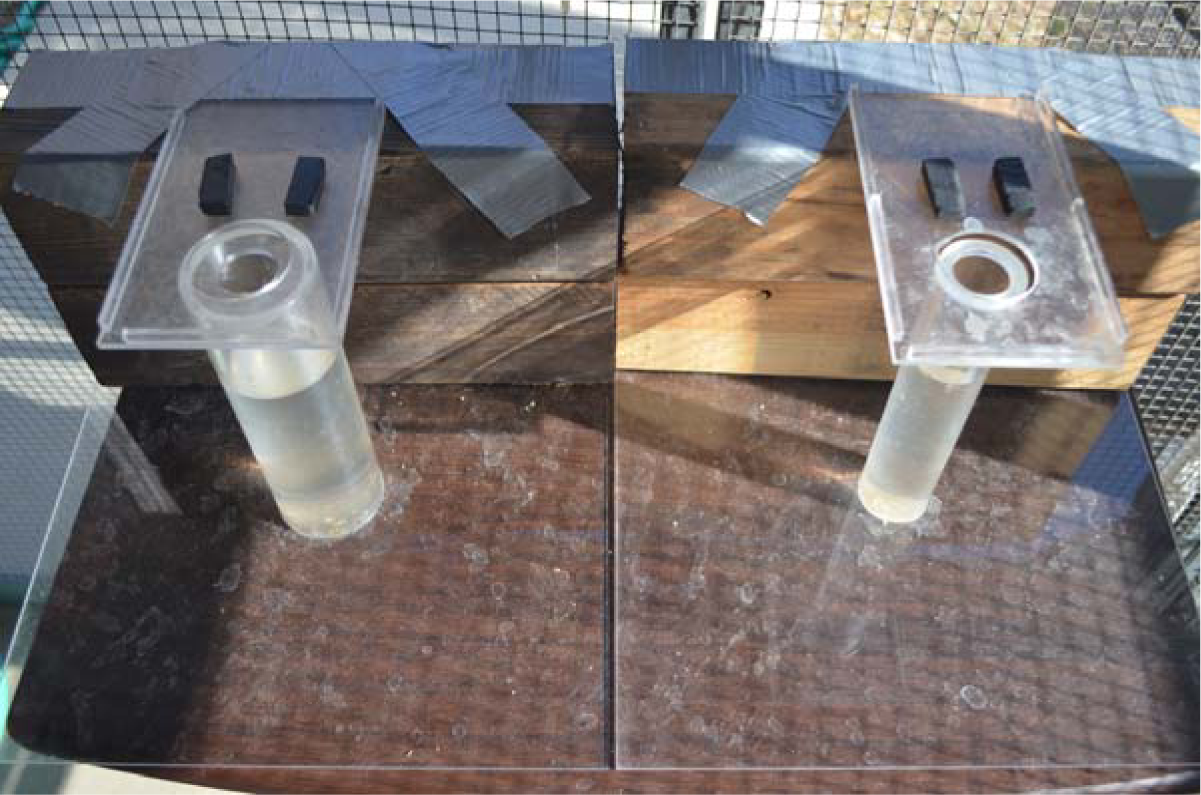
Dropping clay objects into the narrow tube in the Narrow vs. Wide equal water level experiment is the only way to reach the floating food.

### Experiment 5: Narrow vs. Wide Unequal Water Levels

Those grackles that passed Experiment 4 continued to this experiment to determine whether their tube choices adjusted to changing circumstances. This experiment was the same as Experiment 4, except the water level in the narrow tube was lowered to 5cm from the table, thus making the food unreachable even if all objects were dropped into this tube (as in Logan et al. 2014). The water level in the wide tube was raised such that the bird could reach the food in 1-4 object drops, and 20 trials were conducted.

### Experiment 6: Color Association Reversal (learning speed)

The methods were the same as in the Color Association task (Experiment 1), except the food was always placed in the silver tube rather than the gold tube, thus forcing the bird to reverse their preference to consistently obtain the food. Because many other experiments occurred between Experiments 1 and 6, I first checked whether the grackles remembered Experiment 1 before moving them to Experiment 6. If they were successful in 9 or 10 out of their first 10 trials, indicating that they remembered that the food was always in the gold tube, then they moved onto reversal learning with the food always in the silver tube. If they were not successful in their first 10 trials, then they were given a refresher on Experiment 1 until they re-passed the original criterion before moving onto reversal learning.

### Experimenters

I conducted Experiments 2-5, and my research assistants (Luisa Bergeron, Alexis Breen, Michelle Gertsvolf, Christin Palmstrom, and Linnea Palmstrom) and I conducted the stone dropping training and Experiments 1 and 6.

### Statistical Analyses

Two analyses were performed on the color association data (Experiments 1 and 6). First, a bird was considered to pass this test if it chose correctly at least 17 out of the most recent 20 trials (with a minimum of 8 or 9 correct chioces out of 10 on the two most recent sets of 10). Once the bird reached proficiency using this analysis, their individual learning strategy was identified using a contextual, binary multi-armed bandit (see McInerney 2010 for a review). It was contextual in that the subject was only allowed to make one choice per trial, and binary because there were two options on the table, one containing a reward and the other containing no reward. I categorized grackle learning strategies by matching them to the two known approximate solutions of the contextual, binary multi-armed bandit: epsilon-first and epsilon-decreasing (McInerney 2010). The following equations refer to the different phases involved in each strategy:

*Equation 1 (exploration phase)*: 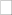 N
*Equation 2 (exploitation phase)*: (1-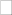) N

N is the number of trials given, and epsilon, 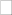, represents the subject’s uncertainty about the location of the reward, starting at complete uncertainty (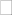=1) at the beginning of the experiment and decreasing rapidly as individuals gain experience with the task and switch to the exploitative phase. Because the grackles needed to learn the rules of the task, they necessarily had an exploration phase. The epsilon-first strategy involves an exploration phase followed by an entirely exploitative phase. The optimal strategy would be to explore each color in the first two trials, and then switch to an exploitative strategy, in which case there would be no pattern in the choices in the exploration phase. In the epsilon-decreasing strategy, birds would start by making some incorrect choices and then increase their choice of gold gradually as their uncertainty decreases until they reach a 100% success rate. In this case, a linear pattern emerges during the exploration phase.

To make the water tube research comparable with previous studies, I used binomial tests to determine whether each grackle chose particular objects or tubes at random chance (null hypothesis: p≥0.05) or significantly above chance (alternative hypothesis: p<0.05). The Bonferroni-Holm correction was applied to p-values within each experiment to correct for an increase in false positive results that could arise from conducting multiple tests on the same dataset.

Generalized linear mixed models (GLMMs) were used to determine whether birds preferred particular objects or tubes (response variable: correct/more correct or incorrect/less correct choice) in a water tube experiment and whether the trial number or bird influenced choices (explanatory variables: experiment, trial number, bird), and to control for the non-independence of multiple choices per trial (random factor: choice number). I used minimal belief priors (V=1, nu=0) and fixed the variance component to one (fix=1) because the measurement error variance was known, as is standard when choices are binary (Hadfield 2010). I ensured that the Markov chain for this test model converged by manipulating the number of iterations (nitt=150000 for the null model, nitt=600000 for the test model), the number of iterations that must occur before samples are stored (burnin=30000), and the intervals the Markov chain stores (thin=300) until successive samples were independent as indicated by low (<0.1) correlations (autocorr function, MCMCglmm package: Hadfield 2014a, b) and there were no trends when visually inspecting the time series of the Markov chain (function: plot(testmodel$Sol); Hadfield 2014a,b). I compared this test model to a null model where I removed all explanatory factors and set it to 1.

I determined whether the test model was likely given the data, relative to the null model by using Akaike weights (range: 0-1, all model weights sum to 1; Akaike 1981; Weights function, MuMIn package: Bates et al. 2011). The Akaike weight indicates the “relative likelihood of the model given the data” (Burnham and Anderson 2002, p. xxiii) and models with Akaike weights greater than 0.9 are considered reliable models because they are highly likely given the data (Burnham and Anderson 2002). The test model was highly likely given the data (Akaike weight=1.00) and the null model was not (Akaike weight=3.4e-30). To investigate potential effects of season or order of testing, I carried out a GLMM to investigate whether the batch to which the bird belonged (explanatory variable: batch=1 or 2) influenced their test performance (response variable: correct or incorrect choice) while controlling for the non-independence of multiple choices per trial (random factor: choice number). The null model was highly likely given the data (Akaike weight=0.94), while the batch model was not (Akaike weight=0.06), indicating that batch did not influence test performance. GLMMs were carried out in R v3.2.1 (R Core Team 2016) using the MCMCglmm function (MCMCglmm package, Hadfield 2014a) with a binomial distribution (called categorical in MCMCglmm) and logit link.

### Data Availability

The data are available at the KNB Data Repository: http://knb.ecoinformatics.org/#view/corina_logan.15.6 (Logan 2015).

## RESULTS

Watch video clips showing examples of each experiment at:http://youtu.be/GhR6fGG1yc4.

### Experiment 1: Color Association (learning speed)

According to the first analysis, all grackles reached criterion in 20-40 trials (Table 1). In the binary multi-armed bandit analysis, Refresco used the epsilon-first strategy because he first explored (i.e., made unsuccessful and/or successful choices) and then exploited (i.e., was successful) every trial thereafter: he explored in his first trial (he failed by choosing silver) and then always chose gold after that (Figure 4). The rest of the grackles used the epsilon-decreasing strategy by exploring more at the beginning and gradually increasing their success until they reached 100% by the end of the experiment (Figure 4). Horchata and Jugo had exceptions to this strategy: Horchata started a second exploration phase at the end of her experiment, and Jugo’s pattern of exploration did not linearly increase at the beginning of his experiment. Jugo did not appear to follow any particular strategies during his learning phase such as ‘always choose the left side’ or ‘always alternate sides’, therefore it is unknown what exploration strategies he used.

**Table 1.**
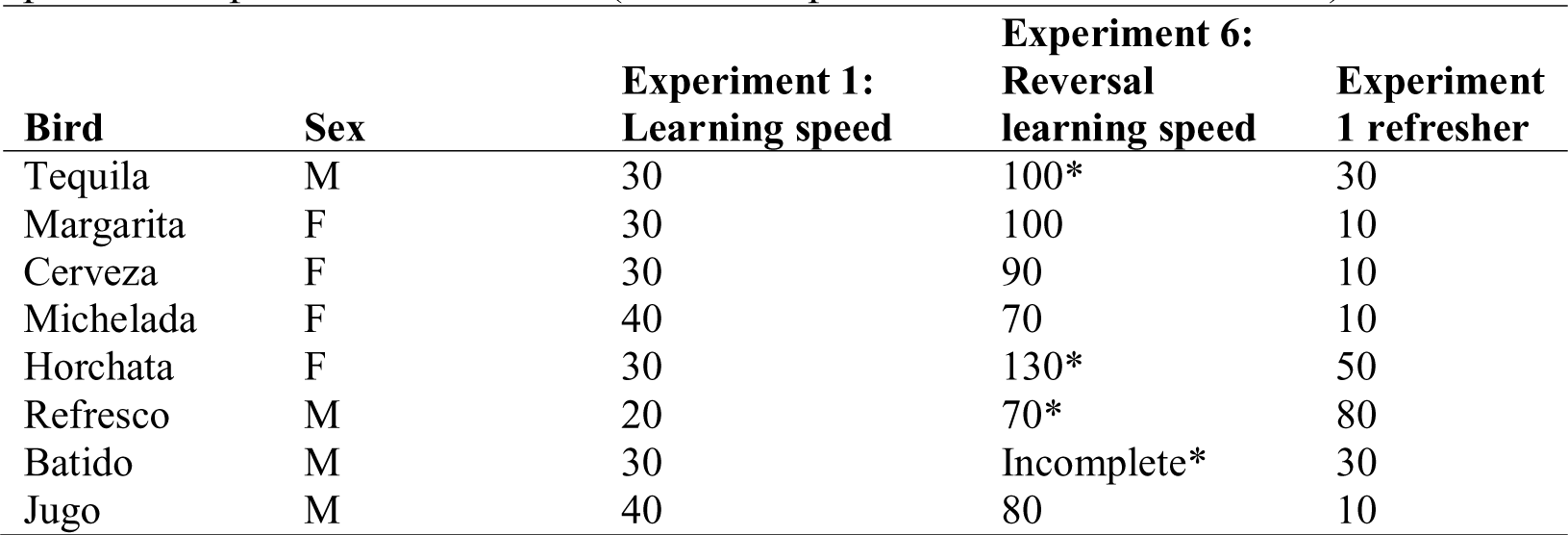
Color association results: The number of trials needed to reach proficiency in Experiment 1 (learning speed), and Experiment 6 (reversal learning speed), and the number of trials needed to pass the Experiment 1 refresher (*=did not pass the refresher in 10 trials).

| Bird | Sex | Experiment 1:<br>Learning speed | Experiment 6:<br>Reversal<br>learning speed | Experiment<br>1 refresher |
| --- | --- | --- | --- | --- |
| Tequila | M | 30 | 100* | 30 |
| Margarita | F | 30 | 100 | 10 |
| Cerveza | F | 30 | 90 | 10 |
| Michelada | F | 40 | 70 | 10 |
| Horchata | F | 30 | 130* | 50 |
| Refresco | M | 20 | 70* | 80 |
| Batido | M | 30 | Incomplete* | 30 |
| Jugo | M | 40 | 80 | 10 |

**Figure 4.**
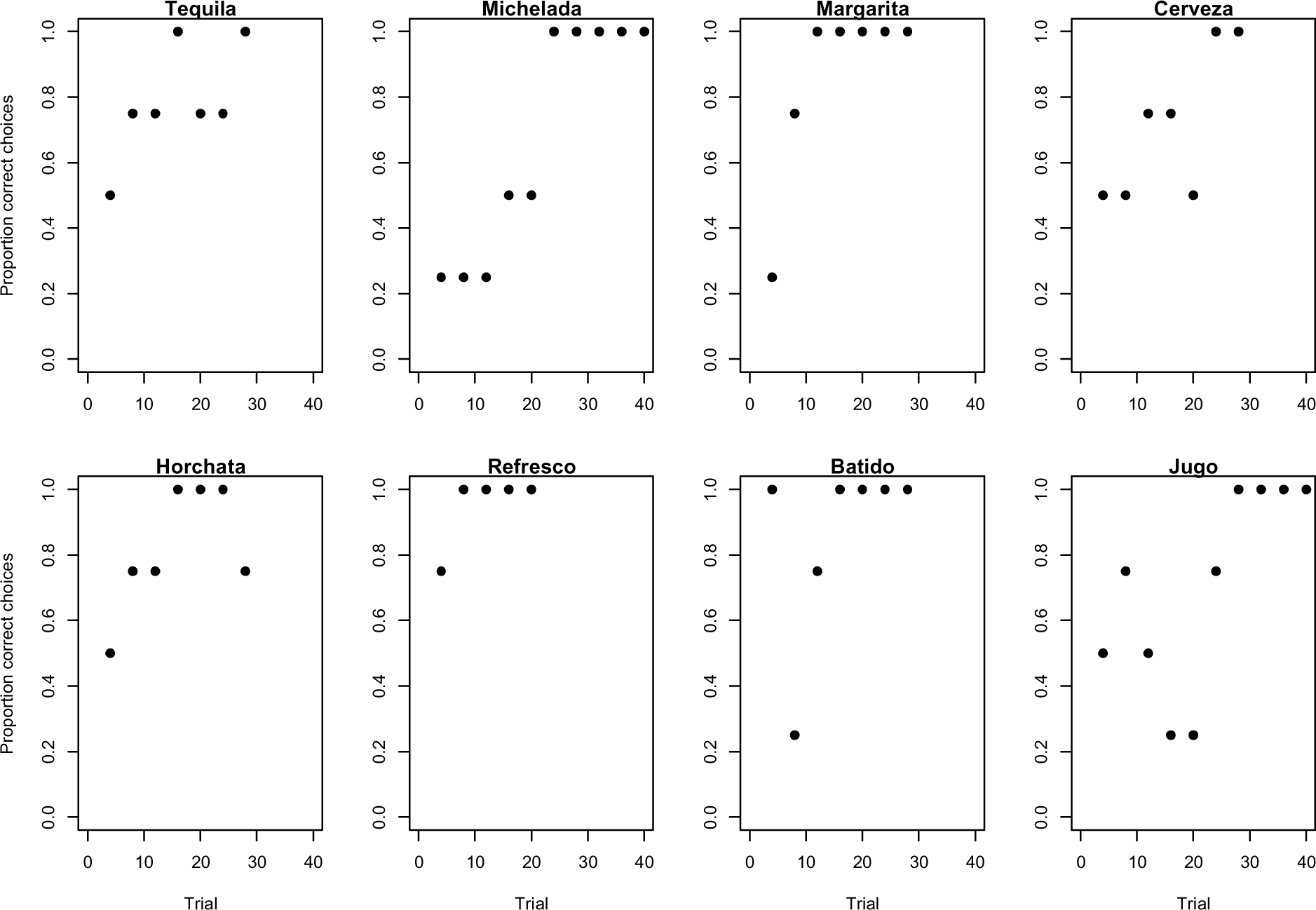
Learning strategies employed by grackles when learning to associate the gold tube with food in Experiment 1 as shown by the proportion of correct choices (non-overlapping sliding window of 4-trial bins) across the number of trials required to reach the criterion of 17/20 correct choices.

### Spontaneous Stone Dropping

No grackle spontaneously dropped stones down the tube of the platform apparatus. Therefore, they all underwent stone dropping training.

### Stone Dropping Training

Most grackles learned to push stones into a tube on the platform apparatus in 135-362 trials (Table 2), however Michelada was scared of the stone falling down the tube and did not habituate to this event and Jugo learned too slowly to become proficient by the time he needed to be released, therefore they were excluded from the stone dropping experiments. The training procedure was modified from Logan et al. (2014) to allow stone pushing from a clear cast acrylic ramp placed on top of the tube rather than stone dropping by picking up the stone from the table and putting it into the tube without a ramp (Figure 1). The modification was necessary because grackles seem to form associations between the stones and the top of the tube, the stones and the table where the food comes out, and the stones falling only in one direction: down. When I placed the stones below the level of the top of the tube to try to train them to pick the stones up and put them in the top of the tube, the grackles took the stones and dropped them off the side of the apparatus or table, often placing them on the table and then looking at where the platform should have fallen open, awaiting the food. Placing the ramp on the water tubes for the experiments was implemented to mitigate this limitation. Once this change was made, it was no longer necessary to train the grackles to pick up and drop the stones because pushing them into the tube sufficed and required less training.

**Table 2.** Performance per bird per experiment: the number of stone dropping training trials needed to reach proficiency, and p-values from Bonferroni-Holm corrected (within experiment) binomial tests for each experiment (- = was not given this experiment). Note: Tequila was the first bird tested and I did not realise until after I trained him to pick up and drop the stones into the tube that I wanted to only train the other birds to push the stones into the tube to save training time. Therefore, the trial numbers for the other birds refer to proficiency to push objects into the tube, not pick up and drop them. Y=yellow, P=purple, B=blue, O=orange, R=red, G=green.

| Bird (color rings) | Sex | Stone drop training trials | Heavy vs. Light | Heavy vs. Light Magic | Wide vs. Narrow Equal |
| --- | --- | --- | --- | --- | --- |
| **Tequila** (YP) | M | 192 push / 233 drop | 0.003 heavy | 0.60 | - |
| **Margarita** (PB) | F | 362 | 0.00001 heavy | 0.02 heavy | - |
| **Cerveza** (BO) | F | 252 | 0.06 | 0.02 heavy | 1.00 |
| **Michelada** (OR) | F | - | - | - | - |
| **Batido** (OP) | M | 179 | 0.002 heavy | 0.02 heavy | 1.00 |
| **Horchata** (GR) | F | 135 | 0.60 | 1.00 | - |
| **Refresco** (PY) | M | 204 | 0.009 heavy | 1.00 | 1.00 |
| **Jugo** (RB) | M | - | - | - | - |

### Water Tube Proficiency Assessment

Most grackles immediately applied their stone dropping skills to a water tube context as indicated by their proficiency on their first trial (Cerveza, Margarita, Refresco, Batido). Horchata was proficient by her second trial. Tequila did initially apply his stone dropping skills to a water tube context, however his order of experiments was different: he went from determining his reachable distance to an experiment involving a water-filled and a sand-filled tube, filled to equal levels. He participated in three trials, but lost motivation and started to give up on participating in stone dropping all together. The water tube proficiency assessment was then developed to remotivate him to participate in subsequent experiments, and the sand vs. water experiment was eliminated. After this additional experience, Tequila needed 76 trials to reach proficiency again.

### Accidental Object Insertions

Because objects were placed near the top of the tube to allow birds to push objects into the tube, it was also possible to accidentally push or kick an object into the tube. Accidental insertions were noted (see Tables 4-6) and included in analyses because birds could learn about the affordances of the task if an object fell into the water, regardless of whether it was chosen or accidental. Some trials were allowed to consist of only an accidental insertion or insertions because the bird was losing motivation and would not have finished the trial otherwise. Counting these as trials errs on the conservative side because throwing the data out and not counting it as a trial removes the ability to account for learning in analyses.

### Experiment 2: Heavy vs. Light

Four grackles (Tequila, Margarita, Batido, and Refresco) were 3.4-5.2 times more likely to choose heavy objects rather than the less functional light objects, while two grackles (Cerveza and Horchata) had no preference (they were 0.6-1.4 times more likely to succeed than fail; see Table 2 for binomial test results and Table 3 for GLMM results). Cerveza and Horchata’s performances improved across trials: they were 3.9-4.4 times more likely to succeed than fail as trial number increased, indicating that they learned through trial and error that the heavy objects were more functional (Table 3). The other grackles’ performances did not improve with increasing trial number, indicating that they used prior knowledge to solve the task (Table 3). Horchata was not motivated to participate in the water tube experiments: she required bait between almost all trials to get her to continue to interact with the apparatus, which might have influenced her lack of success. All choices in all trials for all birds is presented Table 4.

**Table 3.** Examining the influence of experiment, trial, and bird on test success (Test Performance) and whether success increased with trial number (Learning Effects), thus indicating a learningeffect. GLMM: Choices Correct ~ Experiment*Trial*Bird, random = ~Choice Number. CI=credible intervals, italics indicates the intercept.

|  | Test Performance |  |  | Learning Effects |  |  |
| --- | --- | --- | --- | --- | --- | --- |
|  | Posterior Mean | Lower 95% CI | Upper 95% CI | Posterior Mean | Lower 95% CI | Upper 95% CI |
| Choice number | 0.002 | 1.39E-16 | 0.002 | - | - | - |
| *Heavy vs. Light* |  |  |  |  |  |  |
| *Batido* | *1.36* | *-0.03* | *2.83* | *-0.01* | *-0.13* | *0.12* |
| Margarita | 0.29 | -2.31 | 2.83 | 0.04 | -0.17 | 0.26 |
| Cerveza | -1.92 | -4.06 | 0.10 | 0.13 | 0.05 | 0.32 |
| Horchata | -1.01 | -3.14 | 1.14 | 0.01 | -0.20 | 0.19 |
| Refresco | -0.13 | -2.07 | 2.13 | -0.01 | -0.19 | 0.16 |
| Tequila | 0.22 | -1.96 | 2.47 | -0.003 | -0.20 | 0.22 |
| *Heavy vs. Light Magic* |  |  |  |  |  |  |
| Batido | -2.11 | -4.66 | -0.11 | -0.06 | -0.25 | 0.16 |
| Margarita | -0.20 | -3.63 | 3.14 | -0.01 | -0.32 | 0.28 |
| Cerveza | 2.13 | -1.33 | 4.86 | -0.18 | -0.45 | 0.15 |
| Horchata | 2.60 | -0.83 | 5.87 | -0.03 | -0.31 | 0.28 |
| Refresco | -1.32 | -4.73 | 1.92 | 0.33 | 0.03 | 0.62 |
| Tequila | -1.05 | -4.50 | 2.43 | 0.17 | -0.17 | 0.50 |
| *Narrow vs. Wide* |  |  |  |  |  |  |
| Batido | -1.12 | -3.10 | 0.88 | 0.01 | -0.15 | 0.19 |
| Cerveza | -0.21 | -3.61 | 2.74 | 0.06 | -0.20 | 0.33 |
| Refresco | -0.48 | -3.49 | 2.77 | 0.02 | -0.23 | 0.25 |

**Table 4.**
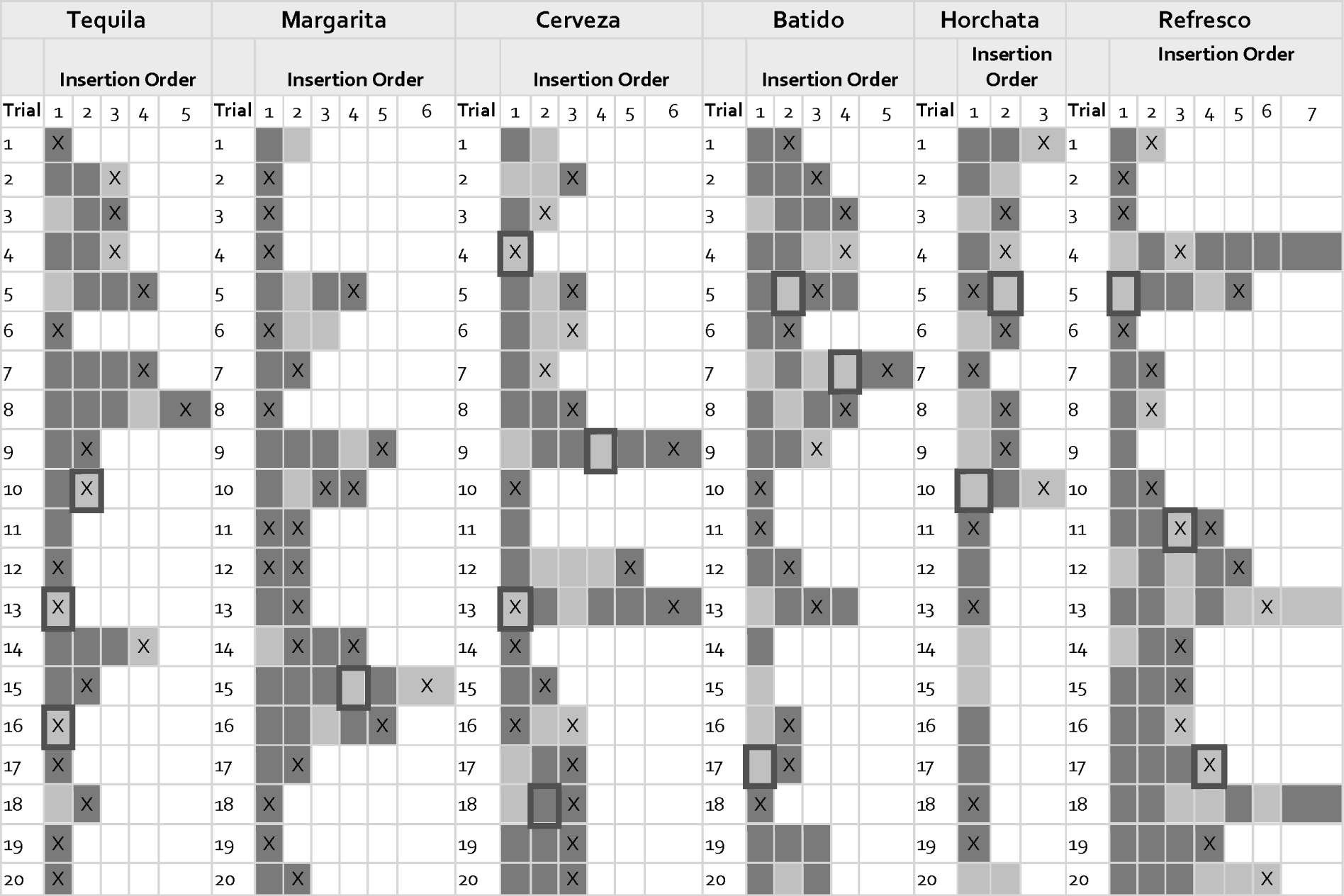
Heavy vs. Light: the order in which the more functional heavy (dark gray) or less functional light (light gray) objects were inserted (columns) into a water tube and whether the bird successfully obtained the food (marked with an X) for trials 1-20 (rows). Choices where objects were accidentally inserted into the tube are indicated by a black border. Accidents were kept in the analyses because they still provided an opportunity for the bird to learn something about the task.

### Experiment 3: Heavy vs. Light Magic

Tequila and Refresco changed from preferring heavy objects in Experiment 1 to having no preference in this experiment, while Batido continued to prefer the non-functional heavy objects (see Table 2 for binomial test results, Table 3 for GLMM results, and Table 5 for all choices made by all birds). Margarita continued to prefer heavy items and Cerveza went from having no preference to preferring the non-functional heavy items because they exhibited an intense interest in the magnet (Table 2; see a video clip at: https://youtu.be/GhR6fGG1yc4). They repeatedly stuck heavy objects to the magnet and attempted to pull them off and required almost no rewards between trials for participating, which indicated a high degree of motivation (motivation that rapidly decreases if they fail experiments). Therefore, I excluded their performances from the behavioral flexibility measure because the experiment did not have the intended effect on their behavior. Tequila gave up after 17 trials, refusing to drop either type of object into the tube, indicating he may have inhibited his choice of heavy. Tequila and Refresco’s performance improved with trial number, indicating that they learned through trial and error about which object was functional (Table 3). The other grackles performances did not change or decreased with increasing trial number, indicating that they did not learn about which object was functional (Table 3). Even though Tequila and Refresco did not learn to prefer light in the amount of trials given, they did exhibit flexibility in that they changed their preferences from heavy in the previous experiment to having no preference in this experiment. Indeed, Refresco would likely have shown a preference for light objects if given more trials because all choices in his last five trials were light objects (Table 5).

**Table 5.**
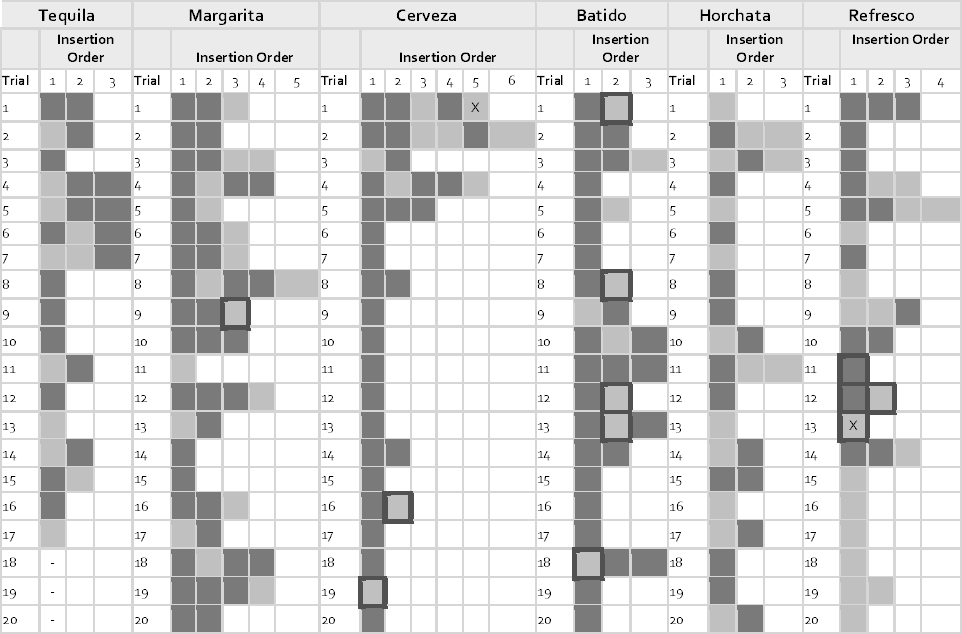
Heavy vs. Light Magic: the order in which non-functional heavy (dark gray) or functional light (light gray) objects were inserted (columns) into a water tube and whether the bird successfully obtained the food (marked with an X) for trials 1-20 (rows). - = did not participate in these trials. Choices where objects were accidentally inserted into the tube are indicated by a black border. Accidents were kept in the analyses because they still provided an opportunity for the bird to learn something about the task.

### Experiment 4: Narrow vs. Wide Equal Water Levels

All three grackles that participated in this experiment displayed no preference for dropping objects into the functional narrow tube or the non-functional wide tube (see Table 2 for binomial test results, Table 3 for GLMM results, and Table 6 for all choices by all birds). None of the grackles’ performances improved with trial number, indicating that they did not learn to distinguish which tube was functional (Table 3). Batido appeared to rely on the strategy of dropping all objects into both tubes regardless of which tube he received a reward from, although in trial 12, he picked up the objects from the wide tube area and dropped them into the narrow tube even though he was only trained to push stones, not drop them (Table 6).

**Table 6.** Narrow vs. Wide Equal Water Levels: the order in which objects were inserted (columns) into functional narrow (dark gray) or non-functional wide (light gray) water tubes and whether the bird successfully obtained the food (marked with an X) for trials 1-20 (rows). Choices where objects wereaccidentally inserted into the tube are indicated by a black border. Accidents were kept in the analyses because they still provided an opportunity for the bird to learn something about the task. Note that Cerveza and Refresco obtained food from the wide tube twice because their motivation changed unexpectedly, thus changing their reachable distance.

|  |  |  |
| --- | --- | --- |
| Cerveza | Batido | Refresco |

|  | Insertion Order |  |  |  |  | Insertion Order |  |  |  |  | Insertion Order |  |  |  |
| --- | --- | --- | --- | --- | --- | --- | --- | --- | --- | --- | --- | --- | --- | --- |
| Trial | 1 | 2 | 3 | 4 | Trial | 1 | 2 | 3 | 4 | Trial | 1 | 2 | 3 | 4 |
| 1 |  |  |  |  | 1 | X |  |  |  | 1 | X |  |  |  |
| 2 |  |  |  |  | 2 |  |  | X |  | 2 |  | X |  |  |
| 3 |  |  |  |  | 3 | X |  |  |  | 3 |  |  |  |  |
| 4 |  |  |  |  | 4 |  |  |  |  | 4 |  |  |  |  |
| 5 |  |  |  |  | 5 |  |  |  |  | 5 |  | X |  |  |
| 6 |  |  |  |  | 6 | X |  |  |  | 6 | X |  |  |  |
| 7 |  |  |  |  | 7 | X |  |  |  | 7 |  |  |  |  |
| 8 |  | X |  |  | 8 |  | X |  |  | 8 | X |  |  |  |
| 9 | X |  |  |  | 9 |  | X |  |  | 9 | X |  |  |  |
| 10 |  |  |  | X | 10 | X |  |  |  | 10 | X |  |  |  |
| 11 |  | X |  |  | 11 | X |  |  |  | 11 |  |  |  |  |
| 12 |  | X |  |  | 12 |  | X | X |  | 12 |  |  |  |  |
| 13 |  |  |  | X | 13 |  | X |  |  | 13 |  | X |  |  |
| 14 |  | X |  |  | 14 | X |  |  |  | 14 |  |  |  |  |
| 15 |  |  |  | X | 15 | X |  |  |  | 15 |  |  | X |  |
| 16 |  | X |  |  | 16 |  |  | X |  | 16 |  |  |  | X |
| 17 |  |  |  | X | 17 | X |  |  |  | 17 |  |  |  |  |
| 18 |  |  |  |  | 18 |  |  | X | X | 18 |  | X |  |  |
| 19 |  | X |  |  | 19 |  | X |  |  | 19 |  |  | X |  |
| 20 |  |  |  |  | 20 |  |  | X |  | 20 |  | X |  |  |

Some grackles did not initially transfer from dropping previous object types to dropping the clay objects used in this experiment. It appeared as though they were trying to solve the problem, but did not perceive the clay objects as being the kind of thing one would drop into a water tube. In these cases, additional training was implemented using a single standard water tube and a mixture of clay objects and stones until the bird was willing to drop objects into the tube even if they only consisted of clay objects. Cerveza transferred to dropping clay objects after 4 training trials, but Tequila and Margarita were excluded from this experiment because they did not transfer to dropping clay objects into tubes. After 14 training trials on a regular water tube with stones and clay objects available to Tequila, it was clear that it would take many more training trials than there was time for and his motivation was greatly diminished. Margarita refused to participate in the training trials. Horchata was also excluded from this experiment because she refused to interact with the objects.

### Experiment 5: Narrow vs. Wide Unequal Water Levels

No grackle passed Experiment 4, indicating they were not sensitive to the differences in water volumes, therefore they were not given Experiment 5, which would have investigated their behavioral flexibility in this context.

### Experiment 6: Color Association Reversal (learning speed)

Margarita, Cerveza, Michelada, and Jugo remembered that food was always in the gold tube because they passed the first 10 trials of the Experiment 1 refresher (Table 1, Figure 5). Tequila, Horchata, Refresco, and Batido needed to re-achieve proficiency on Experiment 1, requiring 30-80 trials before moving onto Experiment 6 (Table 1). Their re-learning patterns followed the epsilon-decreasing strategy that all birds used before, except for Refresco who used the epsilon-first strategy the first time and switched to the epsilon-decreasing strategy for the refresher (Figure 5).

**Figure 5.**
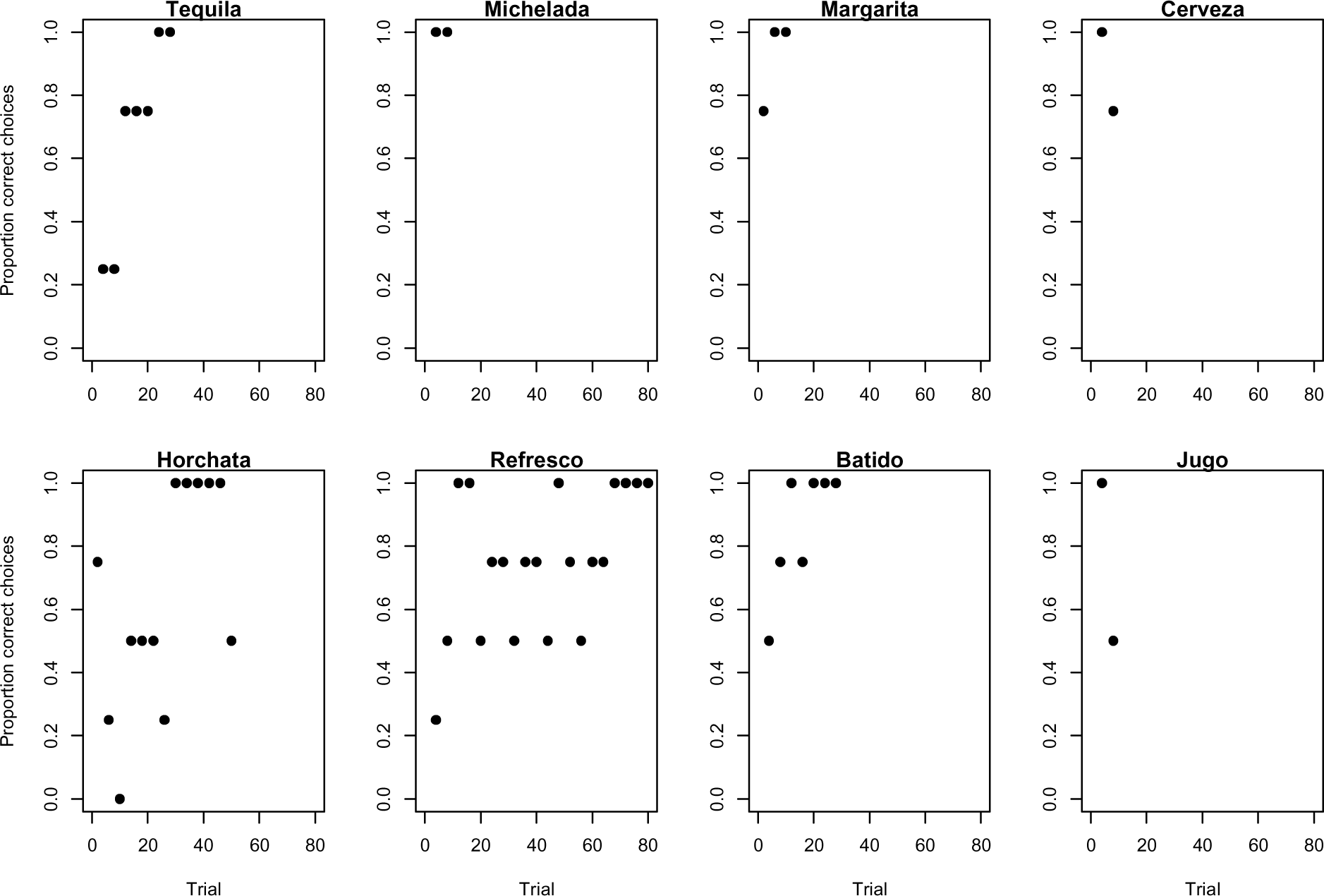
Refresher of Experiment 1: Learning strategies employed by grackles when checking whether they remember that the gold tube contained the food as shown by the proportion of correct choices (non-overlapping sliding window of 4-trial bins) across the number of trials required to reach the criterion of 17/20 correct choices.

Seven out of eight grackles met the reversal learning success criteria (17 out of the most recent 20 trials correct) in 70-130 trials (Table 1), but Batido stopped participating before reaching criterion (Figure 6). All birds used the epsilon-decreasing strategy, but they were slower to learn to reverse their previously learned preference, and many continued to explore throughout the experiment (Figure 6).

**Figure 6.**
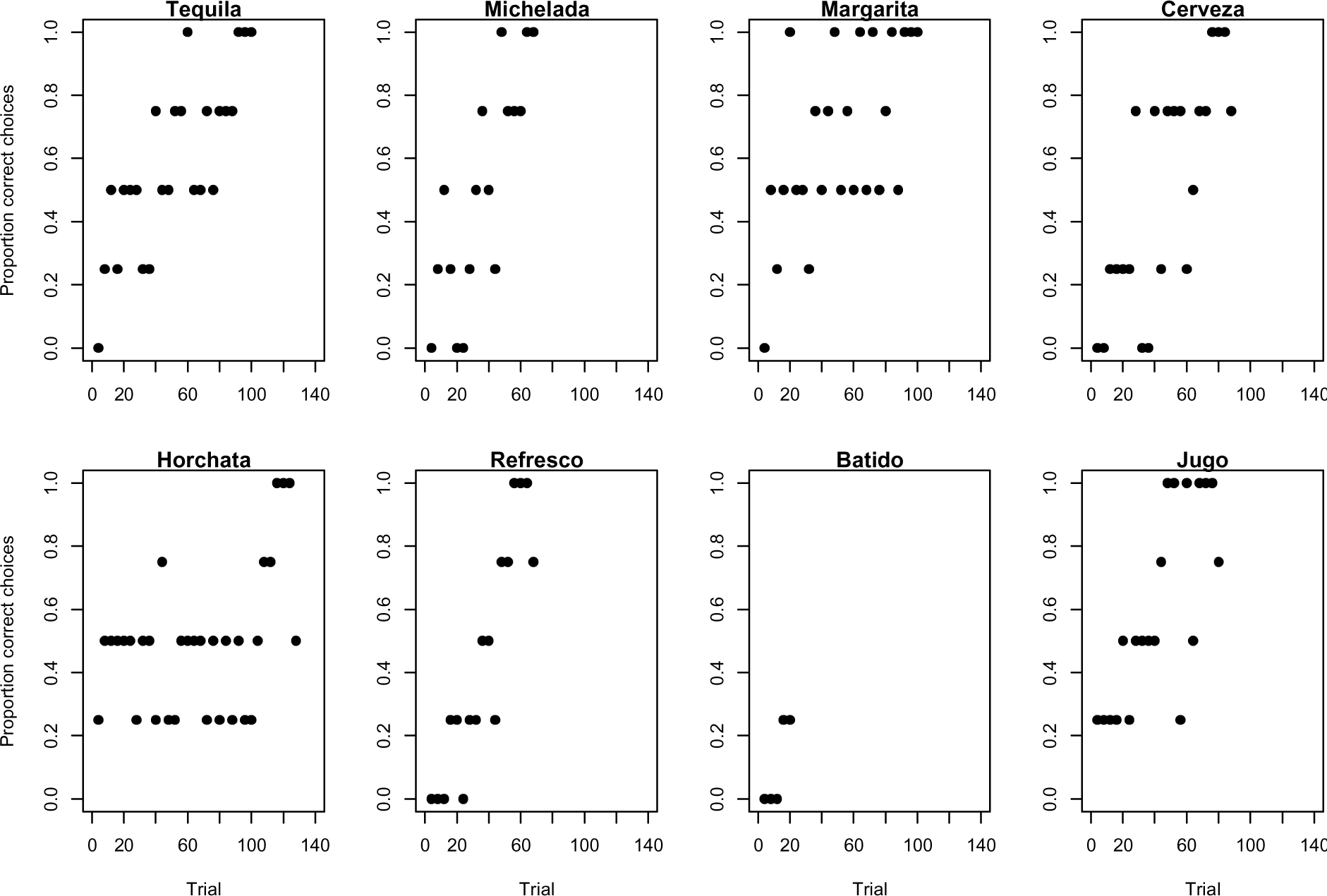
Reversal learning in Experiment 6: Learning strategies employed by grackles when learning to associate the silver tube with food, rather than their previously learned association between the gold tube and food, as shown by the proportion of correct choices (non-overlapping sliding window of 4-trial bins) across the number of trials required to reach the criterion of 17/20 correct choices.

### First Choices on First Trials

All six grackles chose the more functional heavy objects as their first choice in their first trial in Heavy vs. Light, which indicates that they preferred the heavy objects from the very beginning of the experiment (Table 4). Five out of six grackles chose the non-functional heavy objects in Heavy vs. Light Magic (Table 5), which is not surprising given that they had learned to prefer heavy objects in the previous experiment and had likely never interacted with a magnet before, therefore they should have had no reason to have a prior understanding of how the Magic experiment worked. Two out of three grackles chose the functional narrow tube in Narrow vs. Wide with equal water levels, indicating no initial preference for a particular tube (Table 6).

### Did choice number influence the results?

Individuals could learn how the task worked with each choice they made, potentially making each choice dependent on previous choices. Multiple choices could be made per trial; therefore I analyzed how independent choice number was. Choice number was modeled as a random factor in the GLMM and did not influence the results, indicating that choices appear independent of each other (Table 3).

## DISCUSSION

### Grackles are behaviorally flexible and good problem solvers

Despite not being a tool-using species, grackles performed well in the object discrimination tests in this tool-use task. Four out of 6 grackles discriminated between the functional properties of the objects as indicated by their preference for inserting heavy objects significantly more than light objects in the Heavy vs. Light experiment. Their object discrimination performance is similar to that in other successful species where individuals preferred to insert heavy objects which sank rather than light objects that floated and thus were not functional at all: 2/2 Eurasian jays (Cheke et al. 2011), 4/4 New Caledonian crows (Taylor et al. 2011), 6/6 New Caledonian crows (Jelbert et al. 2014), 6/6 New Caledonian crows (Logan et al. 2014), and children age 5 and over (Cheke et al. 2012). This is in contrast to 4-year-old children (Cheke et al. 2012) and Western scrub-jays (Logan et al. 2016) who performed poorly by having no object preference. Perhaps these individuals discriminated between the causal properties of the objects, and thus used causal cognition to solve this task. However, other explanations cannot be ruled out yet: they may have had an innate preference for heavy objects, they might have noticed that inserting heavy objects brings the food closer to the top of the tube than inserting a light object, or they may have associated retrieving food with the heavy objects (Jelbert et al. 2015).

Grackles had a modified version of Heavy vs. Light where the light objects, rather than floating and being non-functional, displaced about half the amount of water as the heavy objects (as in Logan et al. 2016). That most grackles preferred to insert heavy objects when both objects were functional tests a finer degree of object discrimination than has been examined previously, and suggests that they did not simply associate the heavy object with reaching the food because both object types could result in a reward. Three of the 4 grackles that preferred heavy objects did not show a learning effect across the 20 trials in this experiment, indicating that they relied on prior information about the world to solve this task, which suggests that they may have used causal cognition or had an innate preference for heavy objects.

Making heavy and light objects functional in the Heavy vs. Light experiment allowed me to test the object-bias hypothesis, which suggests that individuals solve this experiment because of an innate bias toward heavy objects that are potentially more familiar because they might resemble objects commonly found in the wild (Logan et al. 2014, Jelbert et al. 2015). Accumulating evidence suggests that object-biases are unlikely to be the method by which individuals solve this task: Western scrub-jays (Logan et al. 2016) and 2 grackles had no object preferences in the Heavy vs. Light experiment when both objects were functional. This leaves causal cognition as a likely method for how grackles solved the water tube tasks because they were able to discriminate between the functional properties of the objects, particularly because 2 grackles changed their preference in the Heavy vs. Light Magic experiment, which suggests that they attended to the functionality of the object properties. Western scrub-jays failed to discriminate between object types regardless of their functionality in other Aesop’s Fable tests, therefore it appears that the only reason they passed this experiment is because both objects happened to be functional in this experiment.

Grackles did not discriminate between water volumes in the Narrow vs. Wide equal water level experiment. Perhaps their understanding of water displacement is limited to objects, however more experiments involving object and tube properties would need to be conducted to confirm this.

Grackles were fast to learn an initial preference in the color association task (average 31 trials). Their performance is similar to Western scrub-jays (Logan et al. 2016), 3 species of Darwin's finches (Tebbich et al. 2010), and pigeons (Lissek et al. 2002) who learned in an average of between 40-56 trials. These species are faster than Pinyon jays, Clark’s nutcrackers, a different group of Western scrub-jays (Bond et al. 2007), and Indian mynas (Griffin et al. 2013) who learned on average between 122-280 trials.

Behavioral flexibility was exhibited by grackles because they changed their preferences when the task changed. When the heavy objects in the Heavy vs. Light Magic experiment were no longer functional because they stuck to a magnet, 2 grackles changed from having preferred heavy objects when they were functional in Heavy vs. Light to having no object preference in the Magic experiment. This demonstrates attention to the functional properties of objects in changing circumstances. New Caledonian crows previously showed behavioral flexibility on the Narrow vs. Wide experiments when 4 out of 6 crows preferred to drop objects into the functional narrow tube rather than the non-functional wide tube, and when the wide tube became the functional option, 3 crows changed their preference to the wide tube and 1 changed to no preference (Logan et al. 2014). Grackle performance was similar to the crow that changed from narrow to no preference in the Narrow vs. Wide experiment, although the grackle sample size was reduced from 6 to 4 for the Magic experiment because 2 grackles appeared to be attracted to the magnet and showed a preference for heavy objects as a result. No grackle completely switched their preference to the light objects (as 3 crows did in the Narrow vs. Wide experiments), which may have been due to the difficult design of the apparatus: if one heavy item was inserted, it stuck to the magnet and blocked access to the food regardless of how many light objects were dropped. Thus, grackles had to inhibit inserting any heavy objects to solve this problem, which made the task difficult. Despite the challenging apparatus, Refresco and Tequila likely would have further changed their preference to light objects if given more trials because their performance improved with the number of trials given, indicating that they were learning about the functional properties of the task.

Grackles also demonstrated behavioral flexibility in the color association task by quickly reversing their initially learned preference (average 91 trials). Their performance was similar to 3 species of Darwin’s finches who reversed in an average of 76-95 trials (Tebbich et al. 2010). Darwin's finches and grackles reversed more quickly than pigeons (Lissek et al. 2002), Pinyon jays, Clark’s nutcrackers, Western scrub-jays (Bond et al. 2007), and Indian mynas (Griffin et al. 2013) who learned on average between 142-380 trials.

### Behavioral flexibility varied across contexts

Those grackles that were the most behaviorally flexible in the water tube context (comparing Experiments 2 and 3), were not the most flexible in the color association task (comparing Experiments 1 and 6; Table 7). These results indicate that the context in which behavioral flexibility is tested is important, as suggested by Griffin and Guez (2014). Performing well in these different contexts could require different types of cognition: causal cognition and/or trial and error learning could be used to solve the water tube tasks, while only trial and error learning could be used to solve the color association tasks. Perhaps individuals varied in their reliance on causal cognition, which might have interacted with their reversal learning speed to produce variable results.

**Table 7.**
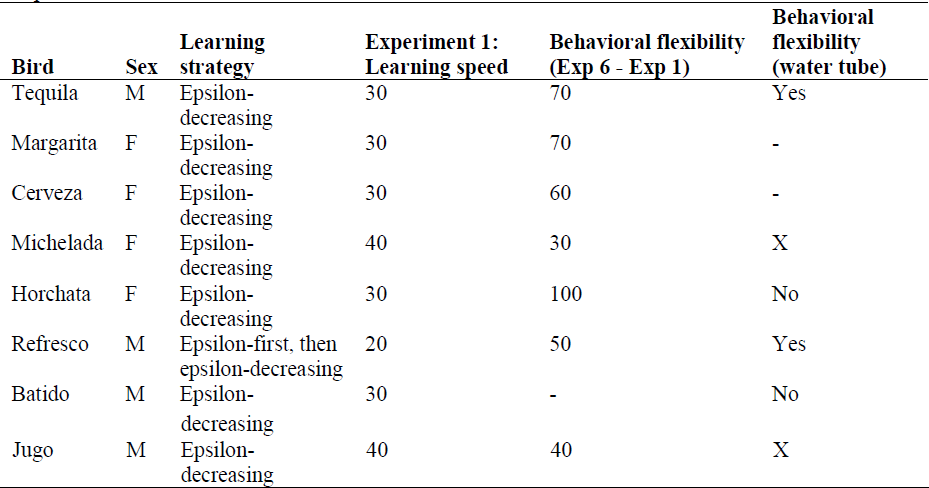
Summary of results: The learning strategy or strategies employed by each bird as well as the number of trials to reach proficiency in Experiment 1, the number of trials needed to reverse their preference (Experiment 6) minus the number of trials needed to initially learn the preference (Experiment 1; a measure of behavioral flexibility), and whether they exhibited a preference change between Experiments 2 and 3. -=preferred to stick heavy objects to the magnet thus the experiment did not test what it was designed to test in these individuals, X=did not participate in this experiment.

### Behavioral flexibility did not correlate with problem solving ability or speed

Grackles that were the most behaviorally flexible were not necessarily the best problem solvers: 4/6 grackles were better problem solvers because they preferred the heavy objects in Heavy vs. Light, however only 2 of these 4 grackles went on to change their preference when the task changed (Table 7). Additionally, those grackles that were the fastest to learn to prefer a color were not the most flexible (i.e., the fastest to reverse this preference) in the color association task. This suggests that behavioral flexibility is an independent source of variation that is distinct from problem solving ability and speed (Cole et al. 2011).

### Those grackles that used more learning strategies were not necessarily more flexible

Refresco was one of the two behaviorally flexible individuals in the water tube experiments, and about average in reversing a color preference (Table 7). He was also the only grackle to use more than one learning strategy in the color association experiment: he used the epsilon-first strategy to sample the environment once before arriving at the correct solution and then he stayed with the correct choice for the rest of Experiment 1. He then switched his learning strategy to epsilon-decreasing for his color learning refresher and for reversal learning (Experiment 6), which is the same strategy the rest of the birds used in both experiments. Individuals using the epsilon-decreasing strategy sample the environment extensively before consistently making the correct choice. Because there was very little individual variation in learning strategies it is difficult to understand how this trait covaries with behavioral flexibility. However, perhaps it is because Refresco was the only one to use multiple learning strategies that he was one of the most behaviorally flexible grackles in one context.

## Conclusion

Results from this investigation demonstrated that individuals differ in behavioral flexibility, which might be a mechanism for maintaining variation within populations – variation that could be useful for successfully adapting to new environments. That behavioral flexibility did not correlate across contexts or with problem solving ability or speed reveals how little we know about behavioral flexibility, and provides an immense opportunity for future research to explore how individuals and species can use behavior to react to changing environments.

## ACKNOWLEDGEMENTS

I am grateful to Luisa Bergeron, Christin Palmstrom, Linnea Palmstrom, and Michelle Gertsvolf for trapping and aviary assistance; Alexis Breen for helping to train Jugo; Brigit Harvey for stone dropping training consultations; Steve Rothstein for scouting grackles and for use of the aviaries; Joe Jablonski and David Bothman for making the apparatuses; Jill Zachary and Kathy Frye at Santa Barbara City Parks and Recreation for use of the Andree Clark Bird Refuge and East Beach Park; Estelle Sandhaus and Chris Briggs at the Santa Barbara Zoo for access to wild grackles; Karrie Black for managing purchasing and grants; Alex Thornton for logistical advice; Dieter Lukas for conceptual input and analysis feedback; Krista Fahy at the Santa Barbara Museum of Natural History for helping with permit applications; Kristine Johnson, Sarah Overington, Julie Morand-Ferron, and Neeltje Boogert for grackle advice and trap plans and manuscript feedback; Manny Garcia for veterinary and permit support; Mary Hunsicker and Bertrand Lemasson for assistance with making the trap; Margaret Tarampi, Eric Egenolf, Rebecca Schaefer, and Sam Franklin for brainstorming object designs; Will Hoppitt for GLMM effect size interpretation assistance; and Irina Mikhalevich and Ljerka Ostojić for manuscript feedback.

## REFERENCES

Akaike H. 1981. Likelihood of a model and information criteria. Journal of Econometrics 16:3–14 doi:10.1016/0304-4076(81)90071-3

Bates D, Maechler M, Bolker B. 2011. lme4: Linear mixed-effects models using S4 classes. Rpackage version 0. 999375–42. http://CRAN.R-project.org/package=lme4. Accessed 12 January 2016.

Bebus, SE, Small TW, Jones, BC, Elderbrock EK, Schoech SJ. 2016. Associative learning is inversely related to reversal learning and varies with nestling corticosterone exposure. Animal Behaviour 111:251–260 doi:10.1016/j.anbehav.2015.10.027

Bird CD, Emery NJ. 2009. Rooks use stones to raise the water level to reach a floating worm. Current Biology 19:1410–1414 doi:10.1016/j.cub.2009.07.033

Bond AB, Kamil AC, Balda RP. 2007. Serial reversal learning and the evolution of behavioural flexibility in three species of North American corvids *(Gymnorhinus cyanocephalus, Nucifraga columbiana, Aphelocoma californica)*. Journal of Comparative Psychology 121:372 doi:10.1037/0735-7036.121.4.372

Buckner C. 2013. A property cluster theory of cognition. Philosophical Psychology 28:307–336 doi:10.1080/09515089.2013.843274

Burnham KP, Anderson DR. 2002. Model selection and multimodel inference: a practical information-theoretic approach, 2nd edn. New York, NY: Springer.

Cheke LG, Bird CD, Clayton NS. 2011. Tool-use and instrumental learning in the Eurasian jay (Garrulus glandarius). Animal Cognition 14:441–455 doi:10.1007/s10071-011-0379-4

Cheke LG, Loissel E, Clayton NS. 2012. How do children solve Aesop’s Fable? PLoS ONE 7:e40574 doi:10.1371/journal.pone.0040574

Chow PKY, Lea SE, Leaver LA. 2016. How practice makes perfect: the role of persistence, flexibility and learning in problem-solving efficiency. Animal Behaviour 112:273–283doi:10.1016/j.anbehav.2015.11.014

Ghahremani DG, Monterosso J, Jentsch JD, Bilder RM, Poldrack RA. 2010. Neural components underlying behavioural flexibility in human reversal learning. Cerebral Cortex 20:1843–1852 doi:10.1093/cercor/bhp247

Griffin AS, Guez D. 2014. Innovation and problem solving: a review of common mechanisms. Behavioural Processes 109:121–134 doi:10.1016/j.beproc.2014.08.027

Griffin AS, Guez D, Lermite F, Patience M. 2013. Tracking changing environments: Innovators are fast, but not flexible learners. PLoS ONE 8:e84907 doi:10.1371/journal.pone.0084907

Hadfield J. 2010. MCMCglmm: Markov chain Monte Carlo methods for generalised linear mixed models. http://citeseerx.ist.psu.edu/viewdoc/download;jsessionid=E395FF7367BF580D916E465F55302C60?doi=10.1.1.160.5098&=rep1&=pdf Accessed 7 May 2015.

Hadfield J. 2014a. MCMCglmm: MCMC generalised linear mixed models. R package. https://cran.r-project.org/web/packages/MCMCglmm/index.html Accessed 30 July 2015.

Hadfield J. 2014b. http://cran.r-project.org/web/packages/MCMCglmm/vignettes/CourseNotes.pdf. MCMCglmm course notes. Accessed 7 May 2015.

Jelbert SA, Taylor AH, Cheke LG, Clayton NS, Gray, RD. 2014. Using the Aesop’s fable paradigm to investigate causal understanding of water displacement by New Caledonian crows. PLoS ONE 9:e92895 doi:10.1371/journal.pone.0092895

Jelbert SA, Taylor AH, Gray RD. 2015. Investigating animal cognition with the Aesop’s Fable paradigm: Current understanding and future directions. Communicative & Integrative Biology 8:e1035846 doi:10.1080/19420889.2015.1035846

Lefebvre L, Whittle P, Lascaris E, Finkelstein A. 1997. Feeding innovations and forebrain size in birds. Animal Behaviour 53:549–560 doi:10.1006/anbe.1996.0330

Lefebvre L, Nicolakakis N, Boire D. 2002. Tools and brains in birds. Behaviour 139:939–973 doi:10.1163/156853902320387918

Lissek S, Diekamp B, Güntürkün O. 2002. Impaired learning of a colour reversal task after NMDA receptor blockade in the pigeon(Columba livia) associative fore-brain (Neostriatum Caudolaterale). Behavioral Neurosciences 116:523–529 doi:10.1037//0735-7044.116.4.523

Logan C. 2015. Great-tailed grackle behavioral flexibility and problem solving experiments, Santa Barbara, CA USA 2014-2015. KNB Data Repository. https://knb.ecoinformatics.org/#view/corina_logan.15.6

Logan CJ, Harvey B, Schlinger BA, Rensel M. 2016. Western scrub-jays do not appear to attend to functionality in Aesop’s Fable experiments. PeerJ 4:e1707 doi:10.7717/peerj.1707

Logan CJ, Jelbert SA, Breen AJ, Gray RD, Taylor AH. 2014. Modifications to the Aesop’s Fable paradigm change performances in New Caledonian crows. PLoS ONE 9:e103049 doi:10.1371/journal.pone.0103049

McInerney RE. 2010. Multi-Armed Bandit Bayesian Decision Making. University of Oxford, Oxford: Technical Report.

Overington SE, Cauchard L, Cote KA, Lefebvre L. 2011. Innovative foraging behaviour in birds: What characterizes an innovator? Behavioural Processes 87:274–285 doi:10.1016/j.beproc.2011.06.002

Peer BD. 2011. Invasion of the emperor’s grackle. Ardeola 58:405–409 doi:10.13157/arla.58.2.2011.405

Pyle P. 2001. Identification Guide to North American Birds Part 1. Ann Arbor, MI: Sheridan Books, Inc.

R Core Team. 2016. R: A language and environment for statistical computing. R Foundation for Statistical Computing, Vienna, Austria. https://www.R-project.org.

Taylor AH, Elliffe DM, Hunt GR, Emery NJ, Clayton NS, Gray RD. 2011. New Caledonian crows learn the functional properties of novel tool types. PLoS ONE 6:e26887 doi:10.1371/journal.pone.0026887

Tebbich S, Sterelny K, Teschke I. 2010. The tale of the finch: Adaptive radiation and behavioural flexibility. Philosophical Transactions of the Royal Society B 365:1099–1109 doi:10.1098/rstb.2009.0291

